# Who’s been sleeping in my bedroom? Sleep and cardiorespiratory physiology when co-sleeping with a dog, cat, or child

**DOI:** 10.64898/2026.06.13.732027

**Authors:** Alexander H.K. Montoye, Dylan Curran, Gregory J. Grosicki

## Abstract

Co-sleeping with pets or children is common, yet its effects on sleep, cardiorespiratory physiology, and behavioral outcomes are understudied. We examined within-person associations between co-sleeping with a dog, cat, or child and sleep characteristics, measures of cardiorespiratory physiology, and next-day physical activity in 1,649,083 person-days from 11,733 adults wearing the WHOOP wearable device. Participants reported nightly co-sleeping via a daily journal in the device’s companion smartphone application, and linear mixed-effects models compared nights with and without co-sleeping within the same individual. Co-sleeping was associated with modest improvements in cardiorespiratory physiology, including lower resting heart rate (0.8-1.1 beats/min), lower respiratory rate (0.04-0.07 breaths/min), and higher heart rate variability (1.41-1.95 ms). Sleep outcomes were mixed, with longer sleep duration (5.5-9.6 min) but more disturbances (0.36-0.40 instances) and slightly less restorative sleep (0.23-1.17%). Associations were generally consistent across groups, although child co-sleeping showed greater sleep disruption. Next-day physical activity was higher following dog and cat co-sleeping (7.6 and 7.4 intensity-weighted min, respectively) but lower following child co-sleeping (2.6 intensity-weighted min). Although effect sizes were small (β range: 0.008–0.045), findings suggest that co-sleeping is associated with a trade-off between modest cardiorespiratory benefits and mild sleep disruption, indicating that co-sleeping decisions may be driven more by personal and contextual factors than by concerns about physiologic impact.

## Introduction

Sleep is essential for overall health and both physical and cognitive function [1], yet a significant proportion of adults in Westernized societies have insufficient or low-quality sleep, with roughly one-third sleeping less than 7 hours per day [2] and nearly ∼30% reporting nighttime sleeping difficulties [3]. With the high prevalence of inadequate sleep, it is important to determine factors that influence sleep quantity and quality. Because sleep occurs within a broader social and environmental context, identifying factors that influence sleep quantity and quality, including sleeping arrangements, is an important area of inquiry. Co-sleeping is common, with 30-50% of adults sharing a bed with a child, pet or both [4, 5]. Despite the common practice of co-sleeping, the physiologic consequences of these arrangements remain incompletely understood.

A recent review by Andre et al. [6] provides a comprehensive overview of co-sleeping research. Co-sleeping with a child shows a mismatch between improved subjective (i.e., self-report) sleep and more disrupted objective (i.e., polysomnography or device-based) sleep measures, especially among mothers. When co-sleeping with animals, species-specific sleep patterns may influence human sleep. Dogs are polyphasic and typically experience several sleep-wake transitions during the night, whereas cats are crepuscular and tend to be more active in the evening and early morning, when humans are more typically asleep [7–9]. Additional factors such as movement and allergen exposure may further contribute to sleep disruption [10, 11], though pets may provide security or other psychological benefits that could improve sleep quality [6, 12]. Empirical studies reflect this balance of competing influences. Hoffman et al. [13] reported that nighttime dog movements were associated with increased movement in human bed partners. Similarly, Patel et al. [14] found poorer sleep quality when dogs shared the bed compared to being present in the room. Although fewer studies have examined cat co-sleeping, available evidence suggests that self-reported sleep disruption may be similar for cat and dog owners [15].

Despite growing interest in co-sleeping, much of the existing literature is limited by small sample sizes, reliance on self-reported sleep outcomes, and between-person designs that compare habitual co-sleepers and non-co-sleepers rather than examining within-person differences. Such designs risk confounding if the two groups differ systematically in demographics, sleep habits, or other lifestyle factors. Moreover, even less is known regarding how co-sleeping relates to cardiorespiratory markers such as resting heart rate (RHR) and heart rate variability (HRV). Poor sleep is associated with elevated RHR and lower HRV [16, 17], suggesting that co-sleeping could influence these measures, but this has not been directly examined at scale.

Large-scale, passively collected wearable data offer new opportunities to address these gaps. Accordingly, we examined within-person associations between co-sleeping with a dog, cat, or child and nightly sleep outcomes, measures of cardiorespiratory physiology, and next-day physical activity in a large sample of wearable device users.

## Materials and methods

### Study design and participants

This study was a retrospective cohort analysis using data from adult users (18-79 yrs) of a wearable fitness tracker (WHOOP Inc., Boston, MA), designed, conducted, and reported in accordance with the Strengthening the Reporting of Observational Studies in Epidemiology (STROBE) guidelines for cohort studies [18]. Participants were randomly selected from the full database if they met predefined inclusion criteria designed to ensure both data completeness, similar between-group demographics, and within-person variability. Eligible users had at least 14 days (and up to 365) of continuous data in 2024, with no missing sleep or biometric measures. Additionally, participants completed daily journal entries in the WHOOP companion smartphone application about the presence of a dog, cat, or child in the bedroom during sleep.

To enable within-person comparisons, participants were required to report the companion (dog, cat, or child) as present on at least three days and absent on at least three days. After the original sample was identified, individuals who appeared in multiple groups (e.g., reported sleeping with a dog and a child) were removed to isolate each individual effect. Prior to matching (described later) the sample included 61,347 individuals (n=44,678 dog co-sleepers, n=12,758 cat co-sleepers, n=3,911 child co-sleepers) contributing 9,966,572 person-days of data. Demographics (age and sex) and anthropometrics (height and weight) were self-reported, and BMI was calculated using the values recorded nearest to the study start date (January 1, 2024). All users consented for their anonymized data to be used for research purposes. The study was reviewed and approved by Salus IRB (Protocol #251121; Austin, TX) and deemed exempt from ongoing oversight based on minimal risk criteria. The de-identified dataset was accessed for research purposes on 4 December 2025. All data were fully anonymized prior to access by the research team, and the authors did not have access to any information that could identify individual participant during or after data collection.

### Wearable device and measures

The WHOOP strap combines accelerometry and photoplethysmography to detect major sleep periods of at least one hour in duration. Within each detected sleep period, WHOOP classified each 30-second interval into one of four sleep stages: wake, light, slow wave sleep (SWS), and rapid eye movement (REM) sleep. Throughout each sleep period, beat-to-beat intervals, along with features derived from frequency transformations, were used to estimate RHR. Additionally, the WHOOP derived peak-to-peak heartbeat intervals from the photoplethysmography signal to estimate HRV, calculated as the root mean square of successive differences (RMSSD). To generate nightly RHR and HRV values, WHOOP first filtered out epochs identified as wake or as having low signal quality. The remaining epochs were then aggregated using a weighted average, in which epochs with higher likelihood of SWS and epochs closer to the end of the sleep were assigned higher weights. Sleep duration was defined as total sleep time (light, SWS, REM). Restorative sleep was defined as the sum of time spent in SWS and REM sleep. WHOOP has been validated against electrocardiography and polysomnography for heart rate (99% agreement) and 2-stage sleep categorization (86-89% agreement) [19, 20].

Daily physical activity was detected automatically using heart rate and motion data from the WHOOP strap or logged manually on the WHOOP smartphone application by the participant. Maximum heart rate was estimated using the non-linear formula 192–0.007*age² [21] manually set, or defined as the highest recorded heart rate during an exercise period with high signal quality. Continuous heart rate data during each activity were used to calculate time spent in five zones defined as percentages of heart rate maximum: zone 1 (50-60%), zone 2 (60-70%), zone 3 (70-80%), zone 4 (80-90%), and zone 5 (90-100%). Minutes in each zone were multiplied by zone-specific weights (i.e., zone 1=1, zone 2=2, etc.), then summed to yield a composite, summated heart rate zone score (SHRZS), which we refer to as physical activity (i.e., intensity-weighted min) [22]. In November 2024, the platform transitioned to heart rate reserve-based zones; as discussed later, we performed a sensitivity analysis using data from prior to this transition (January-October) and found that inclusion of November and December data did not change study outcomes.

### Self-reported co-sleeping with dog, cat, or child

Co-sleep behavior was self-reported via the WHOOP smartphone application’s daily journal feature. Each morning, users were prompted to answer the following yes/no questions regarding the prior night: “Had a dog in the room while sleeping?”, “Had a cat in the room while sleeping?”, and “Slept with a child in your bedroom?”. Users could also skip any question. Only days with explicit “yes” or “no” responses were included in the primary analysis. Journal data were used to identify nights with and without co-sleeping exposure for each group.

### Statistical analysis

All analyses were conducted using day-level physiological and behavioral data for individuals reporting co-sleeping with a dog, cat, or child. Prior to modeling, data were processed to isolate nights with complete sleep, biometric, and journal responses.

To account for differences in demographic and physiological characteristics across the three co-sleeping groups, a matching procedure was applied. Because the child co-sleeping group had the fewest individuals, it served as the reference population. Individuals in the dog and cat groups were matched 1:1 to participants in the child group using nearest-neighbor Mahalanobis distance [23]. Matching variables included age, sex, BMI, and person-level means of RHR, HRV, sleep duration, and physical activity. This yielded three approximately balanced groups of equal size, minimizing the potential effects of these variables on our analysis (**S1 Table**).

The primary outcomes of interest included nocturnal RHR and HRV, respiratory rate (RR), total sleep duration, number of disturbances, a disturbance index expressed as disturbances per hour of sleep, restorative sleep percentage (SWS + REM), and next-day physical activity. For each exposure, nightly deviations (within-person level) and person-mean values (between-person level) were calculated to model chronic and acute associations. Given the study aim of evaluating whether outcomes differed within individuals across co-sleeping and solitary nights, within-person (nightly deviation) effects were considered the primary parameters of interest, while between-person effects were included to account for stable differences in co-sleeping behavior and outcome levels across individuals, thereby isolating associations with night-to-night changes in co-sleeping status within the same individual. Between-person data are shown in tabular form in **S2–S9 Tables** and described in **S10 Text**.

Linear mixed-effects models were fit separately within each matched cohort (dog, cat, child) to estimate context-specific, within-person associations between nights co-sleeping and each outcome, an approach we preferred over a pooled interaction model for its directly interpretable group-specific effects. Fixed effects included both within-person (nightly deviation) and between-person (person-mean) co-sleeping exposure, along with covariates for age, sex, BMI, season, and weekday vs. weekend, plus a random intercept for ID. To assess robustness, we fit two additional model specifications: (Model 2) adding bedtime and nightly deviations in sleep timing, and (Model 3) further adding prior-day physical activity and prior-night sleep duration. Standardized β values from the models were used to determine effect sizes, with 0.10, 0.30, 0.50 used as thresholds for small, medium, and large effects, respectively [24].

In secondary analyses, interaction terms between co-sleeping exposure and sex were added to test for effect modification. The same three model specifications (base, Model 2, Model 3) were applied.

Given that the method of determining heart rate zones (for the physical activity outcome) changed in November, we ran our models both with all 12 months of 2024 data and with only the 10 months from 2024 leading up to the heart rate zone change. There were no differences in outcomes, so we present data from all 12 months. To assess the robustness of primary findings to the handling of nights with missing co-sleeping journal responses, two sensitivity analyses were conducted. Sensitivity analysis A treated null responses as no co-sleeping (null-as-no) but included a null-response indicator as an additional covariate in each model [25]. Sensitivity analysis B applied multiple imputation by chained equations (MICE; m=5 imputations) to estimate effects under a missing-at-random assumption, pooling results using Rubin’s rules [26].

All analyses were conducted in R (version 4.4.2), with statistical significance set at α = 0.05. Model estimates are reported as point estimates with corresponding 95% confidence intervals. Results are presented with primary emphasis on within-person effects, which directly address the study aim, and between-person effects are presented in the **Supporting Information**.

## Results

### Participant characteristics

The final matched dataset consisted of 3,911 unique users per group, for a total sample of 11,733 individuals contributing 1,649,083 person-days (dog: 458,051 co-sleeper days [slept with dog in bedroom on 67.6% of nights included in analysis]; cat: 389,075 co-sleeper days [63.1% of included nights]; child: 162,209 co-sleeper days [45.7% of included nights]). Demographic characteristics are presented in **Table 1**. Average participant ages were within two years across groups. Matching also normalized BMI (within 0.5 kg/m²), RHR (within 0.3 beats/min), and HRV (within 2.4 ms). Sleep time was ∼8 minutes lower in the group with a child, who also had a lower activity score (∼16-20 intensity-weighted minutes). Standardized mean differences (SMDs) between groups were <0.20 for all variables except SHRZS (SMD = 0.30–0.37), indicating good overall balance following matching (**S1 Table**). Mean sleep duration was approximately 7.0 hours across groups (dog: 420 min, cat: 420 min, child: 412 min), near the low end of recommended adult sleep duration [27]. Mean resting heart rate was approximately 61 beats per minute across all three groups, indicative of a relatively fit sample [28].

**Table 1.** Demographics of sample.

|  | **Dog group (n=3,911)** | **Cat group (n=3,911)** | **Child group (n=3,911)** |
| --- | --- | --- | --- |
| **Age (years)** | 37.5 (10.3) | 35.8 (10.3) | 36.6 (6.5) |
| **Body mass index (kg/m<sup>2</sup>)** | 26.6 (4.4) | 26.4 (4.7) | 26.8 (4.8) |
| **RHR (beats/min)** | 61.0 (8.0) | 61.2 (8.1) | 61.3 (8.2) |
| **HRV (ms)** | 52.0 (24.3) | 53.2 (24.9) | 50.8 (22.8) |
| **Sleep time (min)** | 420.0 (38.8) | 420.1 (39.2) | 412.0 (42.5) |
| **Daily physical activity (SHRZS)** | 77.1 (58.7) | 73.2 (57.5) | 56.8 (51.6) |
| **% female** | 35.8% | 35.8% | 35.8% |

Data are shown as mean (standard deviation). RHR: Resting heart rate. HRV: Heart rate variability. SHRZS: Summated heart rate zone score.

### Co-sleeping was associated with favorable cardiorespiratory physiology but more disrupted sleep

In the within-person base models (**Figs 1–3**; **S2 Table**), all eight outcome variables were significantly different on nights where participants reported co-sleeping vs. sleeping alone (p < 0.001): RHR (bpm) decreased in all three groups (dog: -1.14 [95% CI: -1.17, -1.11], cat: -0.94 [95% CI: -0.97, -0.91], child: -0.80 [95% CI: -0.84, -0.76]), whereas HRV (ms) increased (dog: 1.95 [95% CI: 1.86, 2.04], cat: 1.41 [95% CI: 1.32, 1.50], child: 1.83 [95% CI: 1.73, 1.93]). RR (breaths/min) also decreased (dog: -0.072 [95% CI: -0.076, -0.068], cat: -0.066 [95% CI: -0.069, -0.062], child: -0.044 [95% CI: -0.049, -0.040]) and total sleep duration (min) increased in all three groups on co-sleep nights (dog: 9.58 [95% CI: 9.11, 10.04], cat: 7.33 [95% CI: 6.85, 7.82], child: 5.49 [95% CI: 4.91, 6.08]). Disturbances (#) were higher in all three groups (dog: 0.38 [95% CI: 0.36, 0.41], cat: 0.36 [95% CI: 0.34, 0.39], child: 0.40 [95% CI: 0.37, 0.44]), and normalized disturbances (per hour of sleep) similarly increased (dog: 0.019 [95% CI: 0.015, 0.022], cat: 0.023 [95% CI: 0.019, 0.026], child: 0.038 [95% CI: 0.033, 0.042]) on co-sleeping nights. Restorative sleep percentage (%) decreased in all three groups with co-sleeping (dog: -0.35 [95% CI: -0.40, -0.29], cat: -0.23 [95% CI: -0.28, -0.17], child: -1.17 [95% CI: -1.24, -1.11]). Finally, next-day intensity-weighted physical activity increased in the dog and cat groups (dog: 7.56 [95% CI: 6.84, 8.28], cat: 7.35 [95% CI: 6.62, 8.09]) but decreased in the child group (child: -2.57 [95% CI: -3.38, -1.77]) following co-sleeping nights.

**Fig 1.**
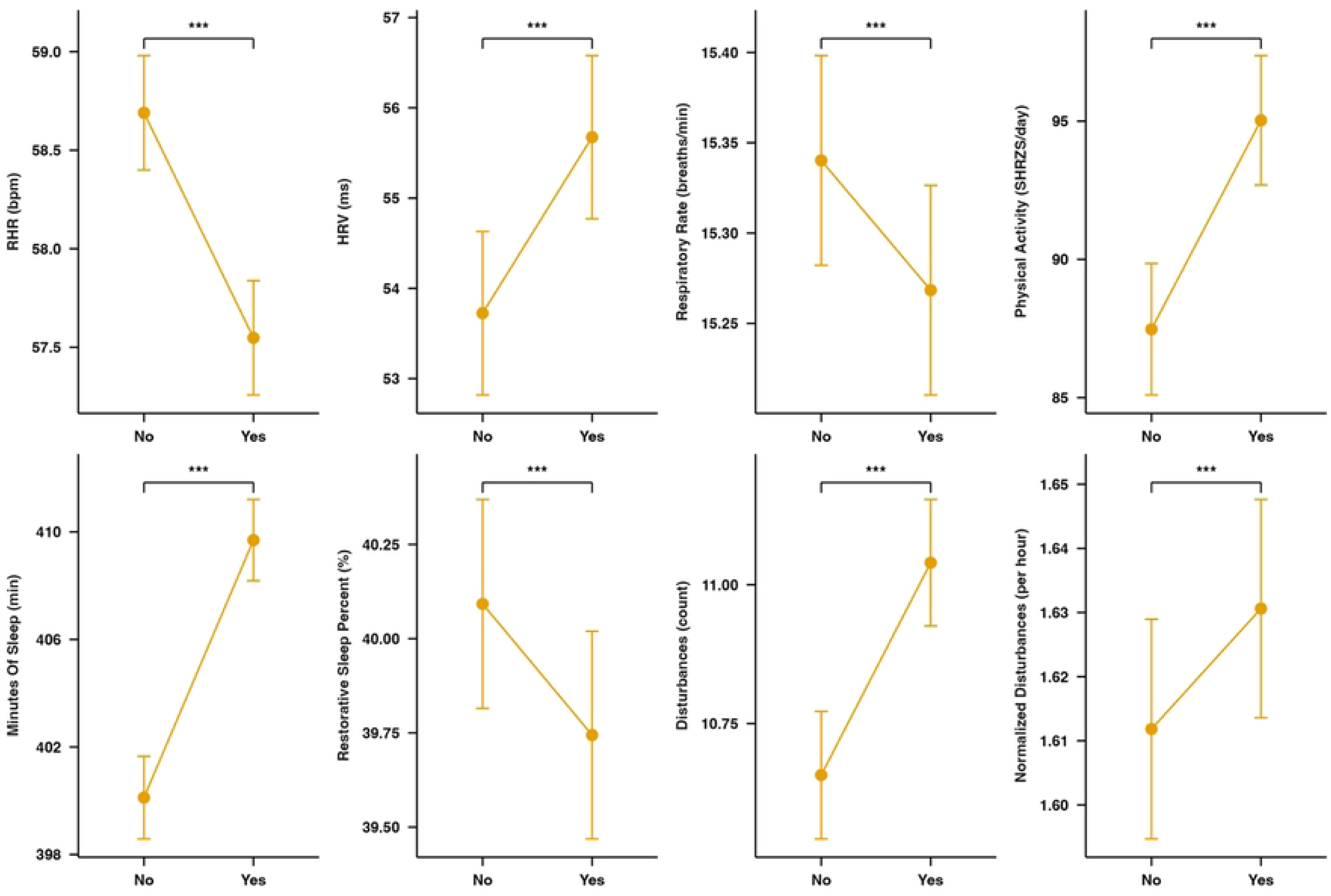
Within-person comparison of co-sleeping vs. not co-sleeping with a dog. RHR: Resting heart rate. HRV: Heart rate variability. SHRZS: Summated heart rate zone score. No: nights not co-sleeping with dog. Yes: nights co-sleeping with dog. ***Indicates significant difference from co-sleeping (p<0.001).

**Fig 2.**
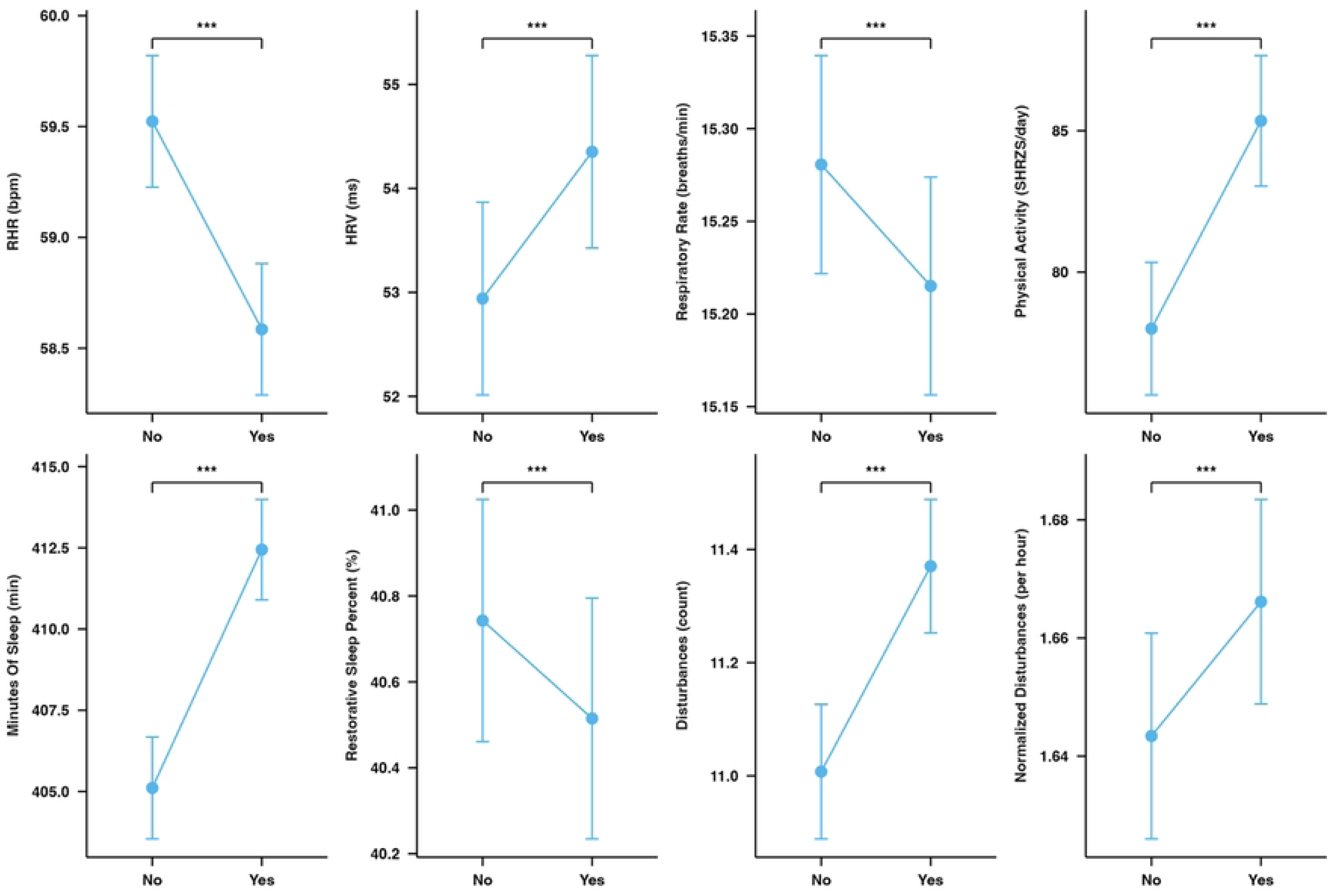
Within-person comparison of co-sleeping vs. not co-sleeping with a cat. RHR: Resting heart rate. HRV: Heart rate variability. SHRZS: Summated heart rate zone score. No: nights not co-sleeping with cat. Yes: nights co-sleeping with cat. ***Indicates significant difference from co-sleeping (p<0.001).

**Fig 3.**
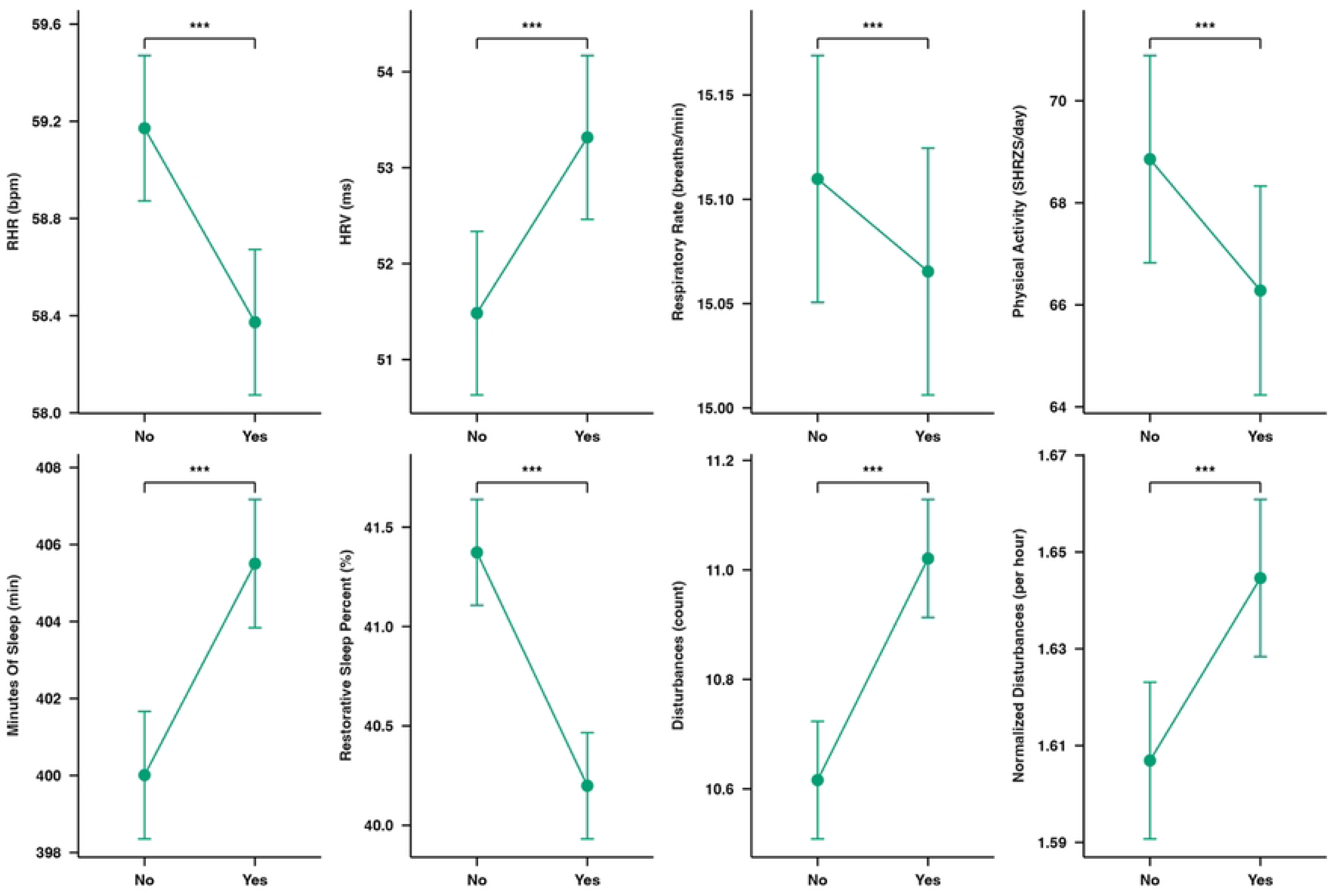
Within-person comparison of co-sleeping vs. not co-sleeping with a child. RHR: Resting heart rate. HRV: Heart rate variability. SHRZS: Summated heart rate zone score. No: nights not co-sleeping with child. Yes: nights co-sleeping with child. ***Indicates significant difference from co-sleeping (p<0.001).

Adjustment for bedtime (Model 2; **S3 Table**) and behavioral covariates (Model 3; **S4 Table**) attenuated the magnitude of several associations but did not change their directionality or significance, with one exception: the within-person association between child co-sleeping and sleep duration became non-significant after full adjustment (+0.4 ± 0.3 min, p=0.106).

### Sex moderated several co-sleeping associations

Co-sleeping effects differed by sex for several outcomes (**S5 Table**). In the dog group, interactions were significant for seven of eight outcomes (all except disturbances, interaction p=0.473): relative to females, males showed larger reductions in RHR and RR, larger increases in HRV and total sleep time, a smaller increase in normalized disturbances, a less pronounced decrease in restorative sleep, and a smaller increase in physical activity (all interaction p≤0.022). In the cat group, interactions were significant for four outcomes (RR, disturbances, normalized disturbances, and restorative sleep; all interaction p≤0.004), with males showing a larger reduction in RR, smaller increases in disturbances and normalized disturbances, and a less pronounced decrease in restorative sleep. In the child group, interactions were significant for five of eight outcomes (RHR, HRV, total sleep time, disturbances, and restorative sleep; all interaction p<0.001): males showed smaller decreases in RHR, smaller increases in HRV, larger increases in total sleep time and disturbances, and a smaller decrease in restorative sleep, whereas no sex differences were observed for RR, normalized disturbances, or physical activity (all interaction p≥0.200).

Adjustment for bedtime (**S6 Table**) and for bedtime plus behavioral covariates (**S7 Table**) attenuated several interactions but did not alter their direction for most outcomes. The most notable exception was child co-sleeping and total sleep time, which decreased in females but increased in males in both the bedtime-adjusted model (females −2.64 vs males +2.12 min; interaction p<0.001) and the fully adjusted model (females −3.00 vs males +2.09 min; interaction p<0.001).

### Findings were robust to sensitivity checks

Two sensitivity analyses were conducted to assess robustness of the primary within-person findings to the handling of nights with missing co-sleeping responses (dog: ∼31%, cat: ∼33%, child: ∼56%). Sensitivity analysis A (**S8 Table**) compared the primary complete-case approach to treating null responses as no co-sleeping (null-as-no) across all groups and outcomes, and the model also included a null-response indicator as a direct covariate alongside the primary model terms across all groups and outcomes. Effect directions and statistical significance were fully preserved across all 24 within-person outcomes, supporting the robustness of primary findings. Sensitivity analysis B (**S9 Table**) applied multiple imputation by chained equations (MICE; m=5 imputations) across all groups and outcomes, finding that all 24 within-person effects remained statistically significant after pooling.

### Effect sizes were small

To assess the practical magnitudes of the observed within-person associations, effect sizes for within-person co-sleeping associations were calculated (absolute value of the standardized β range: 0.008-0.045; data not shown). The largest effects were observed for dog co-sleeping on resting heart rate (β = -0.045) and sleep duration (β = 0.043) and for child co-sleeping on restorative sleep percentage (β = -0.045), with similar effects observed across other outcomes and co-sleeping conditions.

## Discussion

We evaluated the within-person effects of co-sleeping with a dog, cat, or child on sleep, cardiorespiratory physiology, and next-day physical activity using a wearable device. Co-sleeping was associated with modest improvements in cardiorespiratory physiology, including a decreased RHR, increased HRV, and decreased RR concurrent with increased total sleep time but increased disturbances and decreased restorative sleep percentage. These findings were largely similar across co-sleeping groups. Although the combination of improved cardiorespiratory physiology and more disrupted sleep may appear paradoxical, co-sleeping may simultaneously enhance emotional comfort or perceived safety, which has been linked to improved HRV profiles [29–31], while simultaneously introducing movement and disrupting sleep. That the small magnitude of sleep disruption did not appear to offset the cardiorespiratory associations is consistent with these being driven by distinct pathways.

Child co-sleeping showed a more disruptive pattern than pet co-sleeping. Child co-sleeping was associated with smaller increases in sleep duration and greater increases in disturbances and more substantially reduced restorative sleep percentage relative to pet co-sleeping. Between-person effects in the child group were consistent with these trends and persisted after covariate adjustment, suggesting that co-sleeping with children may be less voluntary and more context-dependent than co-sleeping with pets. Prior work has shown that co-sleeping in children is often influenced by caregiving needs, developmental transitions, or family circumstances rather than preference alone [32], and is associated with increased nighttime awakenings, greater caregiver involvement, and poorer sleep indices [33, 34].

Next-day physical activity increased following co-sleeping with a pet (with similar magnitudes in dogs and cats) but decreased in days following co-sleeping with a child. Past research has shown that dog ownership is associated with increased activity levels [35, 36], whereas little research has directly examined cat ownership and physical activity. Decreased activity following co-sleeping with children may reflect fatigue or constrained discretionary time to be active [37, 38]. For example, co-sleeping with a child may limit opportunities for early-morning exercise if caregivers wake simultaneously with the child. These findings suggest that differences in physiologic state and daily time constraints may contribute to the different activity patterns observed following pet vs. child co-sleeping.

Co-sleeping associations varied by sex. Males generally tended to show larger improvements in cardiorespiratory measures and smaller increases in disturbances than females, particularly in the dog group. Most notably, co-sleeping with a child increased sleep time in males but decreased sleep time in females, which may reflect differences in caregiving roles. Previous research has found that mothers are the primary night-time caregivers to young children [39]. However, as with the overall analysis, sex-stratified effects were very small.

Our study has several important strengths including a large sample, a within-person design that reduces confounding from between-person differences, a careful matching procedure that yielded similar demographic and behavioral factors across the three co-sleeping groups, and objective measurement of study outcomes with the wearable device. However, our study also had several caveats worth noting. The Mahalanobis matching resulted in highly comparable groups, although the SMDs do indicate a slightly larger difference in physical activity levels, with the child group less active than the dog and cat groups. Next, co-sleeping was self-reported and a substantial portion of nights had missing journal responses (∼31% for dog, ∼33% for cat, ∼56% for child), though sensitivity analyses confirmed that within-person findings were robust across all outcomes. The WHOOP journal does not distinguish between bed-sharing and room-sharing (e.g., pet was on the floor, or child was in a bassinet), nor does it capture potentially important factors such as bed size, number of co-sleepers, pet size, or child age. Finally, while the WHOOP device provides an objective measure for outcome assessment and has shown good validity for heart rate outcomes (e.g., RHR and HRV) [20, 40], wearable-based measures of sleep architecture, while scalable, do not achieve the accuracy of polysomnography [19, 41, 42].

## Conclusions

Co-sleeping with a pet or child was associated with modest but consistent changes in cardiorespiratory physiology and sleep, reflecting a trade-off between potential calming or comfort-related effects and increased sleep fragmentation. Patterns were broadly similar across companion types, though child co-sleeping showed more pronounced sleep disruption, particularly in females. Effect sizes were small, suggesting that co-sleeping decisions may be guided more by personal factors such as comfort, caregiving needs, and cultural or family routines than by concerns about large impacts on sleep or cardiorespiratory function.

## Acknowledgments

The authors wish to acknowledge the technical assistance of the WHOOP Data Science and Data Engineering teams for their contributions to feature development and data infrastructure. We also thank the broader WHOOP employee community for their continued support of the company’s research initiatives.

## Author contributions

- AHKM: study conceptualization, methodology, formal analysis, visualization, writing – original draft, and writing – review & editing
- DC: software, formal analysis, data curation, visualization writing – original draft, and writing – review & editing
- GJG: study conceptualization, methodology, software, data curation, investigation, resources, formal analysis, project administration, writing – original draft, and writing – review & editing

## AI diligence statement

During the preparation of this work the authors used Claude (Anthropic) for language editing and code review. After using this tool, the authors reviewed and edited the content as needed and take full responsibility for the content of the published article.

## Data sharing

The data and code that support the findings of this study are not publicly available due to intellectual property concerns of WHOOP, Inc. Deidentified participant data and a data dictionary may be made available upon reasonable request to WHOOP (via). Access will require submission of a methodologically sound proposal, approved by WHOOP’s research team, and a signed data use agreement. The timeframe for response to requests will be 4 weeks.

## Supplemental Information

**S1 Table.** Between-group standardized mean differences following Mahalanobis matching.

**S2 Table.** Table of values from unadjusted analysis with within- and between-person effects.

**S3 Table.** Table of values from analysis with within- and between-person effects, adjusted for bedtime.

**S4 Table.** Table of values from analysis with within- and between-person effects, adjusted for bedtime and behavioral covariates.

**S5 Table.** Table of values from unadjusted analysis of sex interaction with within- and between-person effects.

**S6 Table.** Table of values from analysis of sex interaction with within- and between-person effects, adjusted for bedtime.

**S7 Table.** Table of values from analysis of sex interaction with within- and between-person effects, adjusted for bedtime and behavioral covariates.

**S8 Table.** Sensitivity analysis A, where null responses were treated as no co-sleeping but where the model also included a null-response indicator as a direct covariate alongside the primary model terms across all groups and outcomes.

**S9 Table.** Sensitivity analysis B, where multiple imputation by chained equations (MICE) was used to estimate effects under a missing-at-random assumption.

**S10 Text.** Description of between-person effects.

